# Transcriptome profiling reveals gene regulatory network governing seed size in *Brassica napus*

**DOI:** 10.64898/2026.11.06.764911

**Authors:** Yedomon Ange Bovys Zoclanclounon, Laura Siles, Hugh Woolfenden, Richard James Morris, Smita Kurup

**Affiliations:** Rothamsted Research, Harpenden, AL5 2JQ, UK; Department of Computational and Systems Biology, John Innes Centre, Norwich Research Park, NR4, 7UH, UK

**Keywords:** *Brassica napus*, Seed size, Gene regulatory networks, transcriptome analysis, Seed development, polyploidy, crop improvement

## Abstract

Seed size is a critical trait influencing yield and quality in oilseed crops such as *Brassica napus*. Despite their economic importance, the genetic and molecular mechanisms regulating seed size remain unclear, particularly in polyploid species, such as *B. napus*. To deepen our understanding of seed size regulation, we performed a comprehensive transcriptomic analysis across four developmental stages (heart, torpedo, green, and mature) and three distinct seed organs: the seed coat, embryo, and endosperm. Differential expression analysis between *B. napus* var. Zhongshuang (ZS11) and *B. napus* cv. Ragged Jack (RJ) was performed to uncover regulatory networks involved in seed size control. Our results uncovered dynamic gene regulatory networks (GRNs) and pinpointed key regulators, including well-established factors like *ABI5* and *BRI1*, as well as novel candidate genes such as *KNAT5*, *ARF11*, *SUS2*, *UGT72B1*, *UGT89A2*, and *SBT1.7*. The present research provides an understanding of the regulatory networks governing seed size in *B. napus*. These findings not only enhance our knowledge of seed biology but also offer valuable resources for future breeding strategies aimed at optimizing seed size and yield in polyploid oilseed crops.

## Introduction

Seed size is one of the most critical traits from both evolutionary and breeding perspectives, as it directly influences crop yield and quality (Purugganan, 2019; Alam et al. 2022). In oilseed crops, such as *Brassica napus*, *B. rapa*, and *B. juncea*, seed size plays a pivotal role in determining germination success, early seedling vigour, and overall productivity (Liu et al. 2012; Rathnakumar and Sujatha, 2022).

Despite their economic importance, our understanding of the genetic and molecular mechanisms regulating seed size remains limited, particularly in polyploid species like *B. napus,* where subgenome expression bias have been observed (Khan et al. 2022). The A subgenome often dominates during developmental processes like seed development, potentially affecting traits such as seed size. This bias is dynamic and is proposed to be influenced by DNA methylation and small RNAs (Ziegler et al. 2023).

Recent advances in transcriptomic studies have begun to shed light on the regulatory networks underlying seed development in Brassica species (Ziegler et al. 2019; Niu et al. 2020; Dhaka et al. 2022; Khan et al. 2022; Mathur et al. 2022). For instance, in *B. napus*, laser microdissection coupled with RNA sequencing has revealed unique transcriptome profiles associated with specific subregions of the globular seed, including the chalazal endosperm, chalazal proliferating tissue, and chalazal seed coat (Ziegler et al. 2019). These analyses have highlighted the chalazal seed coat’s role in sucrose biosynthesis and transport, while the chalazal endosperm appears to regulate maternal regions through brassinosteroid synthesis (Ziegler et al. 2019).

Extensive research in *Arabidopsis thaliana* and the model crop *Oryza sativa* has elucidated mechanisms underlying seed size regulation (Le et al. 2010; Li and Li, 2015; Li et al. 2019; Ren et al. 2019). Seed development in *A. thaliana* involves stages of embryogenesis, maturation, and consumption of the free nuclear endosperm by expanding cotyledons, followed by rapid desiccation. These processes are controlled by intricate gene regulatory networks (GRNs) involving transcription factors, hormonal signalling pathways, and downstream effectors. For example, the IKU pathway, which includes genes such as *IKU1*, *IKU2*, *MINISEED3 (MINI3)*, and *SHORT HYPOCOTYL UNDER BLUE1 (SHB1)* , regulates early endosperm development and plays a pivotal role in determining final seed size (Li et al. 2019).

In *Brassica rapa*, comparative transcriptome analyses revealed a group of cell cycle-related genes closely linked to seed size variation, with upregulated expression of genes involved in the G2/M cell cycle transition playing a crucial role in promoting seed growth. The transcription factor *Bra.A05TSO1* was shown to positively stimulate the expression of *CYCB1;2* genes, further emphasizing the importance of cell cycle regulation in seed size determination (Niu et al. 2020).

Similarly, in *Brassica juncea,* several DEGs related to cell wall biosynthesis/modification, auxin signalling, cytokinin signalling, and ethylene signalling have emerged as key regulators of seed size (Mathur et al. 2022). For example, genes such as *Dof zinc finger protein DOF3.1* (*BjuVaA01g003707*), *galactosyl transferase family protein XXT3* (*BjuVaA10g002813*), and *endoglucanase 25* (*BjuVaA05g001901*) colocalize with QTLs for thousand-seed weight (TSW) and are candidates for seed size regulation (Mathur et al. 2022). Notably, the upregulation of *C3H TF CYAB1;4* (*BjuVaA01g003080*) in *B. juncea* at early seed developmental stages suggests higher rates of mitotic activity, which potentially could impact seed size.

Despite advances, little is known about the GRNs in *B. napus* across developmental stages. Khan et al. 2022 explored the transcriptome dynamics from gametogenesis to maturation, including morphogenesis, and identified a reduction in seed size associated with *BZIP11* RNAi. Although their study provided valuable insights, grinding full seeds loses organ-specific contributions to seed development. To address this limitation, we designed a comprehensive strategy that covers four key developmental stages (heart, torpedo, green, and mature stages) in three distinct seed organs *viz* the seed coat, embryo, and endosperm. This approach provides insights into both temporal and spatial regulatory mechanisms underlying seed development. Thus, analysing gene expression patterns across stages captures dynamic regulatory networks, revealing key regulators including well-known regulators, *ABI5*, and *BRI1*, and potential new candidate genes, such as *KNAT5*, *ARF11*, *SUS2*, *UGT72B1*, *UGT89A2* and *SBT1.7*.

## Material and methods

### Plant materials, RNA extraction and sequencing

An RNA-Seq time-series study during seed development was conducted in four oilseed rape (OSR) genotypes with different life cycles and environmental requirements: Stellar (spring OSR), Express 617 (winter OSR), Zhongshuang 11 (ZS11, a semi-winter OSR) and Ragged Jack (a heritage kale genotype). Plants were grown under controlled environment conditions with a 16 h photoperiod at 18 °C and 15 °C day and night temperatures respectively, under 400 W HQI lighting (225 µmol m⁻² s⁻¹). For each plant, flowers were manually pollinated and tagged to determine the developmental stage of the embryo. Developing seeds in embryo developmental stage heart, torpedo, green and mature were collected and dissected (Siles et al. 2020) to obtain embryo, endosperm and seed coat for each genotype (**Figure 1a**). High quality RNA was extracted for each tissue type in each seed developmental stage as described in (Siles et al. 2020), and RNA samples were processed by Novogene (Beijing, China). Libraries were sequenced on the Illumina HiSeqX, resulting in 150 base pair paired end reads. The RNA-seq raw data (FASTQ files) from this study (**Table S1**) have been deposited in the European Nucleotide Archive (ENA) with the accession number PRJEB86494 (https://www.ebi.ac.uk/ena/browser/view/ERP169839).

**Figure 1.**
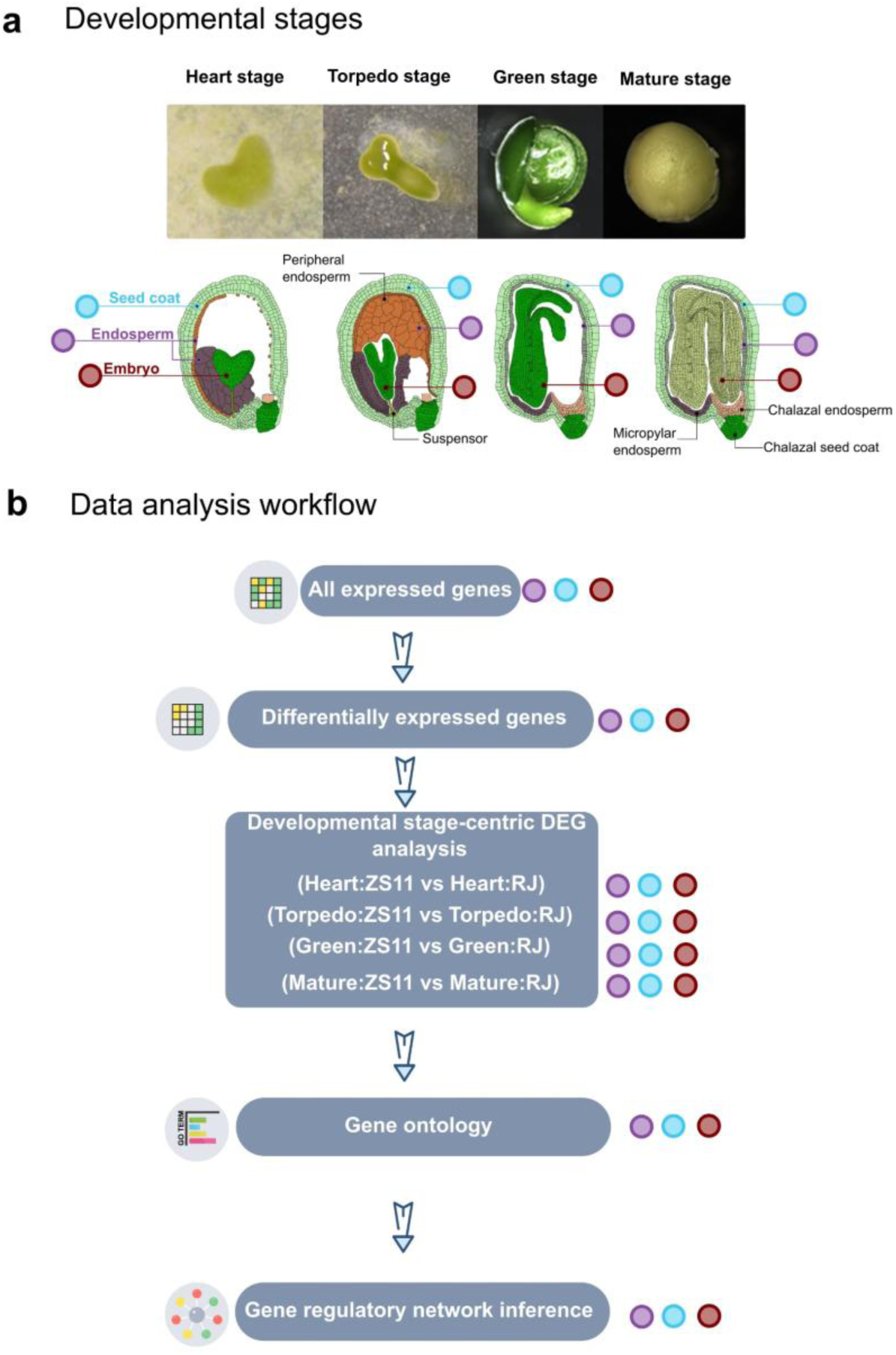
Images and drawings of *Brassica napus* seeds depicting changes in tissue morphology at heart, torpedo, green, and mature embryo stages. (a). RNA was extracted in three tissues: seed coat (light blue circle), endosperm (purple circle), embryo (maroon circle). Overview of the gene discovery analysis workflow showing the gene expression analysis steps from the expression count matrix to network inference (b). Images in (a) were adapted from (Siles et al. 2020).

### Phenotyping assay and seed area data analysis

Once the plants were fully dry and mature, pods were manually collected and opened to obtain the seeds. Seeds were pooled from different plants for each genotype. As per genotype, a total of 300 seeds were imaged (using a NIKON D5300 with NIKKOR 50 mm f/1.8 prime lens, Minato, Tokyo, Japan) on a photobench, and seed area (mm²) was measured with the image analysis software Fiji (Schindelin et al. 2012).

The seed area data were analysed in R v4.2.2 (R Core Team, 2022). Normality within each genotype was assessed using the Shapiro-Wilk test implemented in shapiro.test() from the stats v3.6.2 package. Homogeneity of variances was evaluated using Levene’s test via leveneTest() in the car v3.1-3 package (Fox and Weisberg, 2019). The data deviated significantly from a normal distribution (Shapiro-Wilk, p < 0.05) and showed unequal variances among genotypes (p < 0.05). Consequently, differences in seed area among genotypes were examined using Welch’s one-way ANOVA followed by Games-Howell post-hoc test, implemented with ggbetweenstats() in the ggstatsplot v0.13.3 package (Patil, 2021).

### Transcriptome analysis

RNA raw data were quality checked with FastQC v0.11.9 (Andrews, 2010) and pre-processed with Trimmomatic v0.39-2 (Bolger et al. 2014) for adapter removal and quality trimming. The clean data were then mapped onto *B. napus* Darmor v4.1 reference genome (Chalhoub et al. 2014) using HISAT2 v2.1.0 (Kim et al. 2015) with the settings “–pen-noncansplice 20 –mp 1,0” (Baruzzo et al. 2017). Subsequently, reads counting was performed using StringTie v 2.1.1 (Pertea et al. 2016).

Differential expression analyses were conducted in R v4.2.1 using the DESeq2 package (Love et al. 2014).

To gain a detailed insight into transcriptome expression at each developmental stage within each organ, a species-pairwise comparison was performed at each stage for per tissue (**Figure 1b**). The factors species and developmental stage were merged into a single composite factor (“Species_Stage”), allowing the extraction of relevant contrasts, including ZS11_Heart vs RJ_Heart, ZS11_Torpedo vs RJ_Torpedo, ZS11_Green vs RJ_Green, and ZS11_Mature vs RJ_Mature. Genes were considered differentially expressed if |log2 fold change| ≥ 2 with an adjusted p-value < 0.05.

To assess the functional attributes of differentially expressed genes, gene set enrichment and motif analyses were performed with SeqEnrich (Becker et al. 2017). In addition, the top 1000 DESeq2-ranked genes were extracted for downstream co-expression and gene regulatory network analyses.

### Multivariate co-expressed gene profiles clustering

Co-expression analysis of the expression profiles was conducted with the coseq R package (Rau and Maugis-Rabusseau, 2018) embedded in DIANE package v1.0.6 (Cassan et al. 2021). Normalized expression data were arcsin-transformed, and the Gaussian mixture models were fit to each cluster. To evaluate the quality of the clustering, the Integrated Complete Likelihood criterion was employed. Subsequently, the optimal number of clusters was determined via the elbow method, wherein the number of clusters was selected at the point where the Integrated Complete Likelihood exhibited a plateau, indicating minimal marginal gains from further increasing the number of clusters.

### Transcription factor data retrieval

Prior to the gene regulatory network inference, gene regulators (**Table S2**) were defined based on *B. napus* var. Darmor transcription factors retrieved from the Plant transcription Factor Database (PlantTFDB) (Tian et al. 2020) at https://planttfdb.gao-lab.org/index.php?sp=Bna (accessed December 12, 2023).

### Gene regulatory network inference

To reconstruct transcriptional regulatory networks, we employed the GENIE3 (Gene Network Inference with Ensemble of trees) algorithm (Huynh-Thu et al. 2010) along with permutation-based approach implemented in DIANE R package v1.0.6 (Cassan et al. 2021) for network sparsification.

GENIE3 is one of the top-performing methods that quantifies regulatory influences between transcription factors and target genes using a random forest regression framework (Marbach et al. 2012). It calculates an importance metric ranking putative regulator-target links based on the regulator’s predictive strength for the target’s expression pattern across the random forest trees.

In the present study, the regulator-target gene links were initially inferred using the GENIE3 algorithm with a total of 1000 random forest trees. High scoring regulator-target relationships were then selected to construct a network with a connectivity density threshold of 0.03.

To assess the statistical significance of the inferred links, null distributions of importance scores were estimated through permutations of the expression data. The observed GENIE3 importance values were subsequently tested against these null distributions. Regulator-target interactions with importance scores corresponding to a false discovery rate-adjusted p-value below 0.05 were retained in the final gene regulatory network model.

This approach combines GENIE3’s strength in scoring potential regulatory relationships with a rigorous statistical filtering step from DIANE v1.0.6, to enrich for regulatory interactions underlying observed transcriptional changes. The resulting high-confidence gene regulatory network was rendered using Gephi v0.10.1 (Bastian et al. 2009).

## Results

### Seed size varies between *Brassica napus* genotypes with gene expression profiles clustering by developmental stage

We first sought to quantify seed size across four cultivars of *B. napus* cv Stellar DH (Stellar), *B. napus* var. Zhongshuang 11 (ZS11), *B. napus* cv. Express 617 (Express), and *B. napus* cv. Ragged Jack (RJ). Seed area measurements of 300 seeds revealed variability ranging from 0.98 mm^2^ in Express to 5.79 mm^2^ in ZS11 (**Figure 2a**). Among the genotypes, RJ exhibited smaller seed area with an average of 2.14 ± 0.28 mm^2^, while ZS11 had a mean seed area of 3.78 ± 0.55 mm^2^, approximately 1.8 times larger than RJ (p-value < 0.001, Games-Howell post-hoc test following Welch’s one-way ANOVA) (**Figure 2b**).

**Figure 2.**
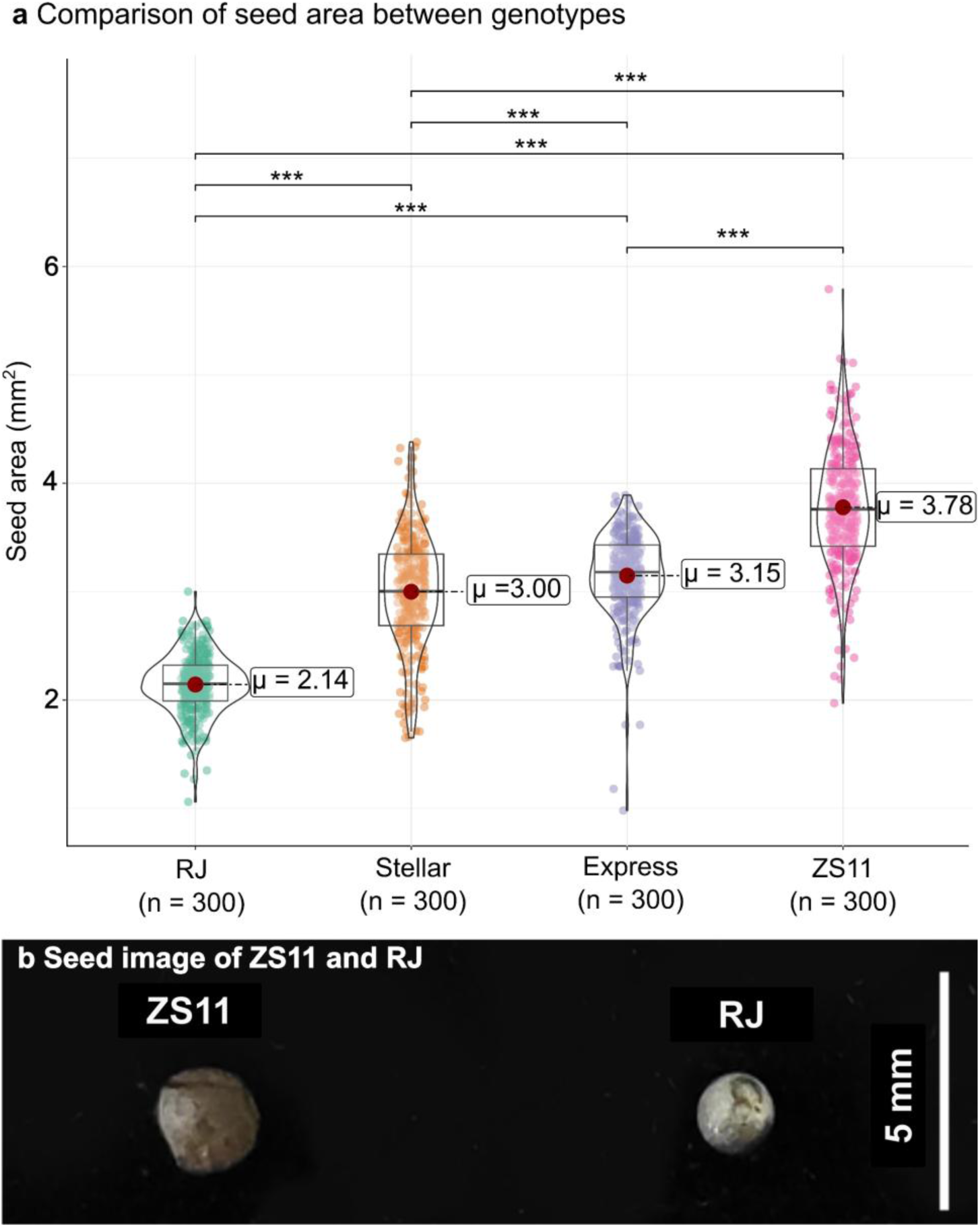
Seed size variability in *B. napus* genotypes. (a) Boxplot showing the data distribution of seed area for the *B. napus* genotypes. Significant pairwise comparisons were highlighted with *** (p < 0.001). (b) Photographs of *B. napus* var. Zhongshuang 11 (ZS11), and *B. napus* var. Ragged Jack (RJ) seeds.

To identify genes associated with these differences in seed size, we selected the two extreme genotypes ZS11 and RJ for downstream comparative transcriptomics at the heart, torpedo, green and mature seed developmental stages.

Gene expression-based clustering revealed a clear distinction of the expression profile by developmental stage (**Figure 3**) between these two genotypes. Within each stage, distinct grouping by tissue was noticeable. Furthermore, the gene expression clustering also delineated the expression profiles by genotypes, highlighting the robustness of the contrast between the ZS11 and RJ genotypes.

**Figure 3.**
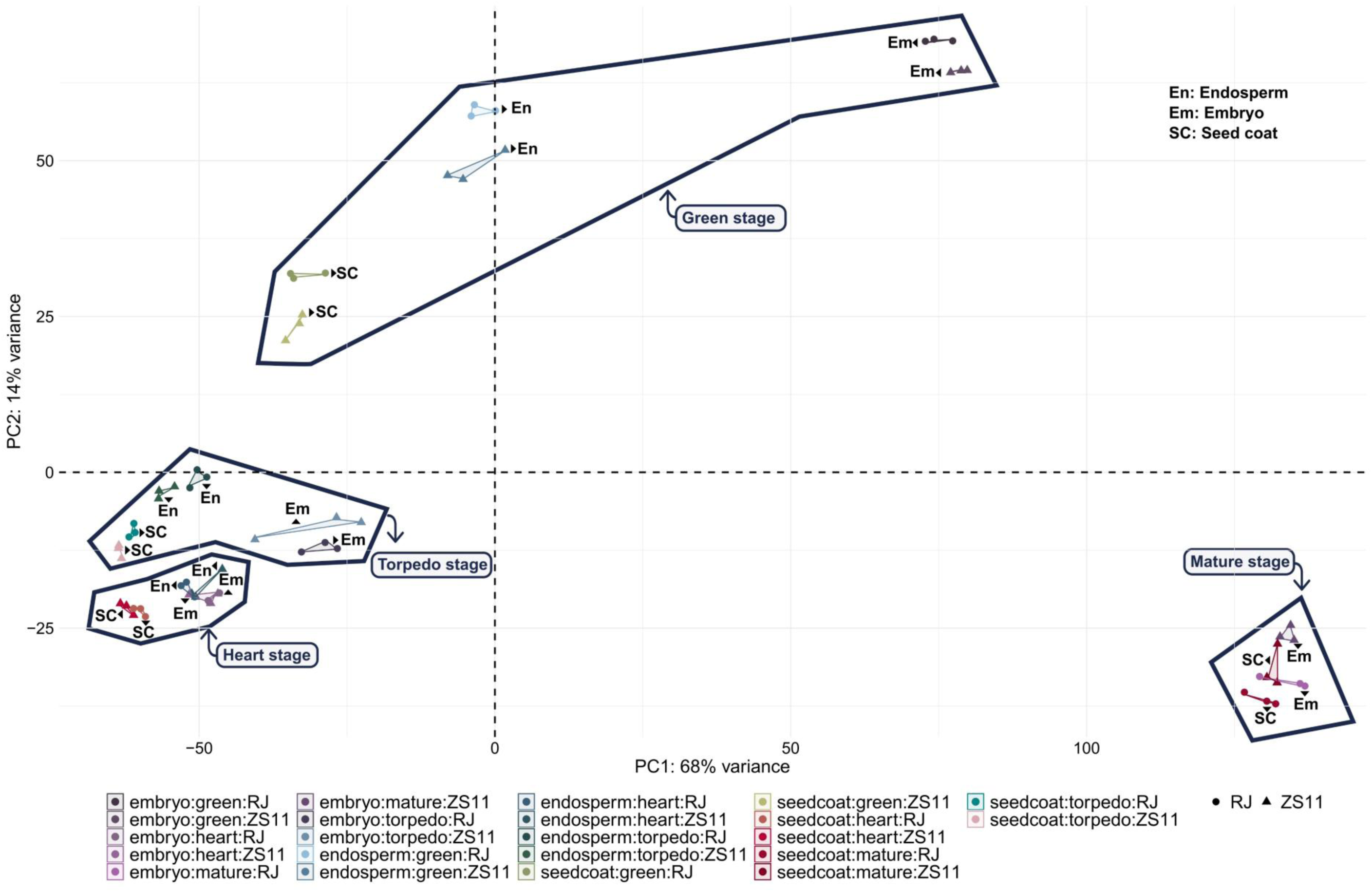
Principal component analysis plot showing the gene expression-based clustering pattern of the samples. Circle symbol represents the small size seed genotype Ragged Jack (RJ). Triangle symbol represents the big size genotype Zhongshuang 11 (ZS11). Symbol colouring was applied to represent three replicates for each tissue type and developmental stage.

### Dynamic transcriptome profiling identifies regulatory genes during different stages of seed size development

To identify candidate regulatory genes for seed size, we performed stage-centric differential expression analysis for each tissue, endosperm (**Figure 4a-d**), embryo (**Figure S1**), and seed coat (**Figure S2**) for both genotypes (ZS11 and RJ).

**Figure 4.**
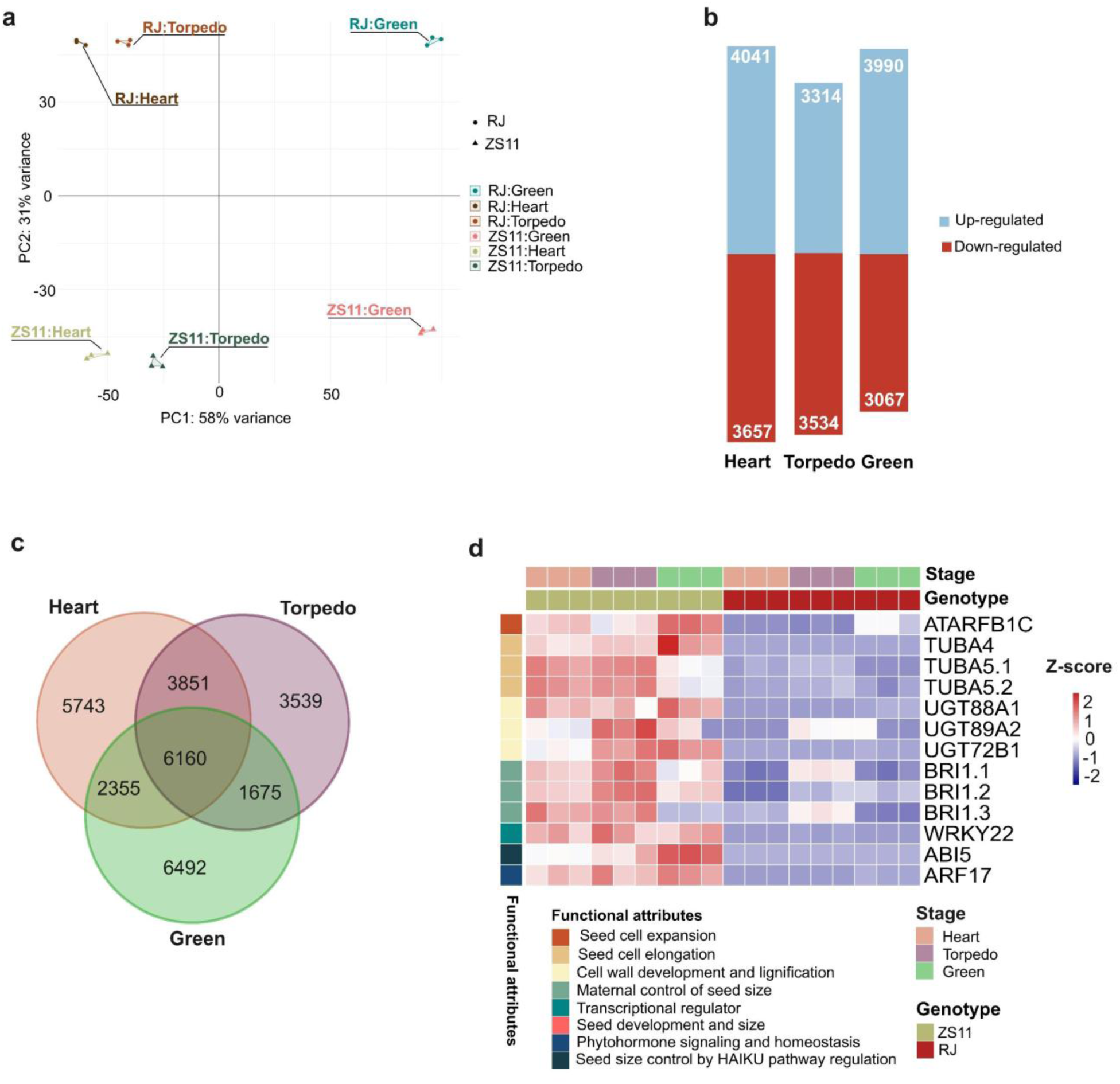
Transcriptome profiling uncovers seed size-related genes. (a) Principal component analysis of the normalized expression profiles in endosperm tissues across developmental stages (heart, torpedo, and green). Circle symbol represents the small size seed genotype Ragged Jack (RJ). Triangle symbol represents the big size genotype Zhongshuang 11 (ZS11). Different colours represent specific endosperm development stage for each genotype. (b) Number of up-regulated (blue colour) and down-regulated (red colour) genes at the heart, torpedo and green stages, respectively. (c) Venn diagram depicting the shared and specific gene number in three contrasting conditions: ZS11 vs RJ at heart, ZS11 vs RJ at torpedo, and ZS11 vs RJ at green stages, respectively. Genes are considered significantly differentially expressed with statistical significance (adjusted p-value < 0.05) and substantial changes in expression (|log2 fold change| ≥ 2). (d) Heatmap of seed size development-related genes identified in endosperm tissues from gene ontology analysis, preferentially expressed in ZS11.

In the endosperm dataset, clustering based on gene expression (**Figure 4a**) highlighted differences not only at the genotype level but also at the developmental stage level. Comparing ZS11 and RJ, a total of 4041, 3314, and 3990 genes were upregulated at the heart, torpedo, and green stages, respectively (adjusted p-value < 0.05, |log2 fold change| > 2) (**Figure 4b**). The number of stage-specific transcripts decreased from 5743 at the heart stage to 3539 at the torpedo stage and increased to 6492 at the green stage (**Figure 4c**).

Examination of the gene attributes via gene ontology (**Table S3**) identified genes associated with seed development (**Figure 4d**). Notably, we identified genes involved in phytohormone signalling and homeostasis pathways regulating seed size development, including three *BRASSINOSTEROID INSENSITIVE 1* (*BRI1*) genes (*BnaC07g47240D*, *BnaA08g16520D*, and *BnaA01g05490D*), which were preferentially highly expressed in ZS11. Among the identified genes, *BnaC04g52290D*, which encodes *ABSCISIC ACID-INSENSITIVE 5* (*ABI5*) and participates in the HAIKU pathway, was also highlighted. The DEG analysis revealed UDP-glycosyltransferase genes, including *UGT89A2* (*BnaC04g44310D*), *UGT72B1* (*BnaA09g00220D*), and *UGT88A1* (*BnaC05g36720D*) (**Figure 4d**), which are associated with cell wall development and lignification. Additionally, genes related to seed cell elongation, such as those encoding alpha tubulin (*TUBA5*, *BnaC02g09010D*, and *BnaC02g09000D*), were detected, and all were highly expressed in ZS11 in comparison with RJ. Interestingly, key regulators belonging to the Auxin response factor (*ARF*) gene family (*ARF17*, *BnaA07g33740D*), and *WRKY* gene family (*WRKY22*, *BnaA09g00120D*) were revealed (**Figure 4d**).

The analysis of embryo-centric transcript-based clustering revealed stage- and species-specific differentiation (**Figure S1a**). Notably, the expression of up-regulated genes varied across stages (**Figure S1b**). Specifically, at the early seed developmental stage (referred to as heart stage hereafter), 2664 genes were upregulated. Subsequently, an increase in gene expression was observed at the middle stages, with 3288 genes at torpedo stage and 3962 genes at green stage. Conversely, at the end of the developmental stage (mature stage), a lower accumulation of gene transcripts occurred, with 2529 genes being upregulated (**Figure S1b)**.

Furthermore, the gene ontology analysis (**Figure S1c**) of the top 1000 differentially expressed genes revealed seed-stage-specific gene expression related to embryo development ending dormancy (GO:0009793), embryo development (GO:0009790), regulation of embryonic development (GO:0045995), and regulation of embryo sac egg cell differentiation (GO:0045694) (**Figure S2c**). Among these seed-stage-specific genes, *BnaA09g52110D*, *BnaC09g02300D*, *BnaA02g05460D*, *BnaC04g02050D*, and *BnaC09g04440D* represent *MATERNAL EFFECT EMBRYO ARREST 47* (*MEE47*), *EMBRYO DEFECTIVE 1276* (*EMB1276*), *EMBRYO DEFECTIVE 2758* (*EMB2758*), *EMBRYO DEFECTIVE 2654* (*EMB 2654*), and *EMBRYO DEFECTIVE 2024* (*EMB2024*) respectively (**Figure S4c**). Interestingly, the gene *BnaA05g22260D*, also known as Nucleostemin-like 1 (*NSN1*), is a GTPase that plays a crucial role in embryogenesis (Wang et al. 2012) was also detected. Moreover, a trehalose-6-phosphate synthase enzyme (*TPS1*) gene (*BnaA07g34230D*), which catalyzes the first step in trehalose biosynthesis and is essential for embryo maturation, along with the *PXG1* gene (*BnaC01g18950D*) involved in oxylipin metabolism and lipid body biogenesis in seeds, and the Ribosome Recycling Factor (*RRF*) gene (*BnaC04g20850D*) facilitating proper translation and protein synthesis required for early embryo development, were pinpointed, and as well as in the endosperm, these genes were more expressed in ZS11 than RJ.

Clustering of gene expression profiles from the seed coat revealed similar patterns, depicting a grouping based on developmental stage and genotypes (**Figure S2a**). Notably, the upregulated transcripts exhibited a stagnated expression, reaching a plateau during the early (3660 at heart stage) and middle developmental stages (3560 at torpedo stage and 3677 at green stage), followed by a drastic drop (1909) of about 1.5 times at the maturation stage (**Figure S2b**). Gene ontology analysis identified genes involved in embryo development ending in seed dormancy (GO:0009783), seed maturation (GO:0010431), seed development (GO:0048316), and regulation of seed maturation (GO:2000034), among others (**Figure S2c**). The analysis of transcript expression highlighted the preferential expression of the cell growth regulator gene family, Basic helix-loop-helix family transcription factor, *BnaC09g00090D* (*bZIP10*), in ZS11 (**Figure S2c**). Additionally, the ABA-related genes *BnaA02g01510D* and *BnaA07g39250D*, encoding for a GEM-like protein 5 (*GER5*) and 9-cis-epoxycarotenoid dioxygenase (*NCED9*) enzyme, respectively, were identified. Notably, the accumulation of embryo-specific genes essential to the process of embryo development was observed. These genes include *BnaA04g02190D* (*EMB1637*), *BnaC03g09230D* (*EMB3013*), *BnaC04g01050D* (*EMB1989*), *BnaC09g04440D* (*EMB2024*), and *BnaA03g09860D* (*MEE70*) (**Figure S2d**). Furthermore, we identified two representatives of the Protein Disulfide Isomerase-Like Protein 1-1 family (*PDIL1-1*), *BnaC04g51940D* and *BnaA07g10870D*, involved in regulating seed protein amount and composition, impacting endosperm development.

Overall, the expression analyses revealed significant transcript accumulation in seed size development in the endosperm tissue compared with the seed coat and embryo, where basal genes for embryo maintenance and cell growth were found to be differentially expressed (**Figure 4, Figure S1, Figure S2**). Subsequently, for co-expression and network analyses, we focussed on the endosperm dataset.

### Co-expression analysis of the endosperm transcriptome reveals candidate gene clusters for the regulation of seed size

To reveal genes associated with developmental stage in the endosperm, we analysed transcript dynamics across stages in ZS11 and RJ. The top 1000 differentially expressed genes partitioned into nine clusters based on their co-expression (**Figure 5, Table S4**). Globally, in clusters 2, 3, 5, and 8, over 400 genes were up-regulated in ZS11 and down-regulated in RJ at the heart, torpedo, and green stages. In cluster 5, genes representing key regulators belonging to the *ARF* (*BnaA07g33740D*, *ARF17*) and *WRKY* (*BnaA09g00120D*, *WRKY22*) gene families, along with genes involved in cell wall development and lignification (*BnaA09g00220D, UGT72B1*, and *BnaC05g36720D*, *UGT88A1*), were found to be co-expressed. A total of 96 genes in this cluster exhibited a stable expression change in ZS11 within the developmental stages, while being barely expressed in RJ. The cluster 8 encompassed 93 genes, including regulators influencing endosperm growth (*BnaC04g52290D*, *ABI5*) and seed cell elongation (*BnaC02g39910D*, *TUBA4*). In ZS11, gene expression in this group steadily declined across developmental stages, whereas in RJ it was marginally expressed. Conversely, the 90 genes in cluster 2, containing the cell elongation gene *BnaC04g44310D* (*UGT89A2)*, exhibited an increasing transcript expression pattern from the heart to the green stage, passing through the torpedo stage in both genotypes with higher expression values in ZS11. Finally, in the cluster 3, showing the largest number (150) of co-expressed genes, a significant stage-specific expression changes were noted with an increase in transcript expression from the heart to the torpedo stage, followed by a drop at the green stage in the ZS11 genotype. Notably, this cluster also contained three copies of the key endosperm growth regulator *BRASSINOSTEROID INSENSITIVE 1* (*BRI1*; *BnaA01g05490D, BnaC07g47240D, BnaA08g16520D*) together with genes involved in seed cell elongation and expansion (*BnaC02g09010*, *BnaC02g09000D*, *TUBA5*, *BnaA05g32370D*, *ARFB1C*).

**Figure 5.**
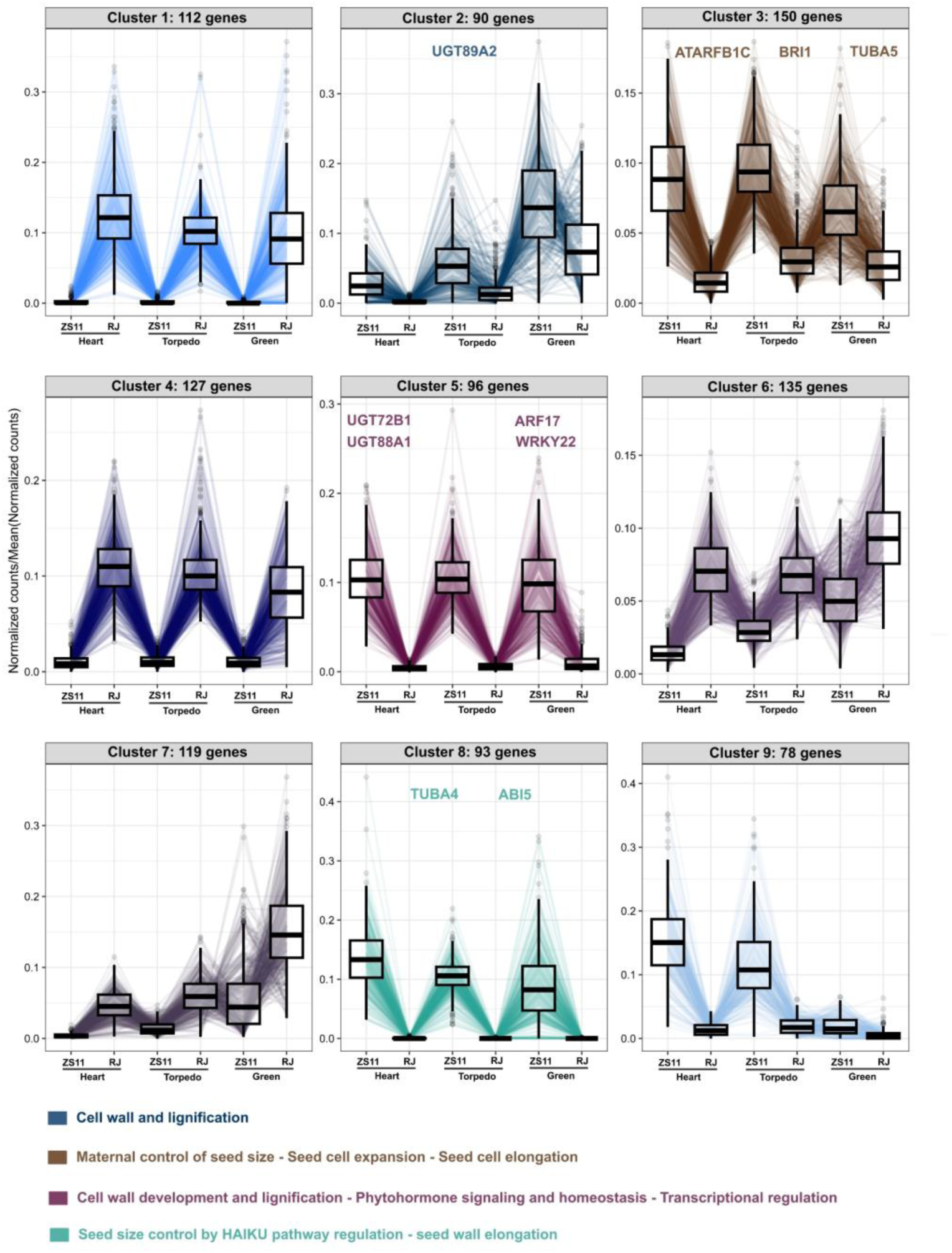
Clustering analysis of the top 1000 differentially expressed genes in endosperm tissues by developmental stage (heart, torpedo, green). Expression profiles represent the DESeq2 normalised expression divided by the mean normalised value across all stages. Genes were clustered into nine groups. Genes involved in seed size development are highlighted at the top of the box plots. Clusters without highlighted genes do not contain key genes related to seed size development. The functional attributes of the genes within each group were described at the bottom of the graph. In each cluster, the circles, whiskers and thick black horizontal line represent the expression of each gene, the first quartiles and the median, respectively. ZS11 stands for Zhongshuang 11, and RJ stands for Ragged Jack respectively.

Based on this co-expression analysis, we identified candidate genes and trends of the transcript kinetics underlying seed growth in the large seed genotype ZS11, along with the developmental stages.

### Inferred gene regulatory network reveals regulator-target genes contributing to seed size

Leveraging the list of transcription factors (**Table S2**) and the endosperm expression profile dataset (**Table S5**), we conducted a gene regulatory inference using random forest regression (Huynh-Thu et al. 2010) in conjunction with permutation statistics (Cassan et al. 2021) to identify candidate regulators and their target genes.

The resulting network generated five communities of interconnected genes, exhibiting a modular and non-assortative architecture, characteristic of biological networks (**Figure 6a**). To minimize bias in the prediction, we grouped together regulators that exhibited high levels of correlation (Spearman correlation threshold r = 0.9) (Cassan et al. 2021), thereby reducing the incrementation of spurious nodes and edges.

**Figure 6.**
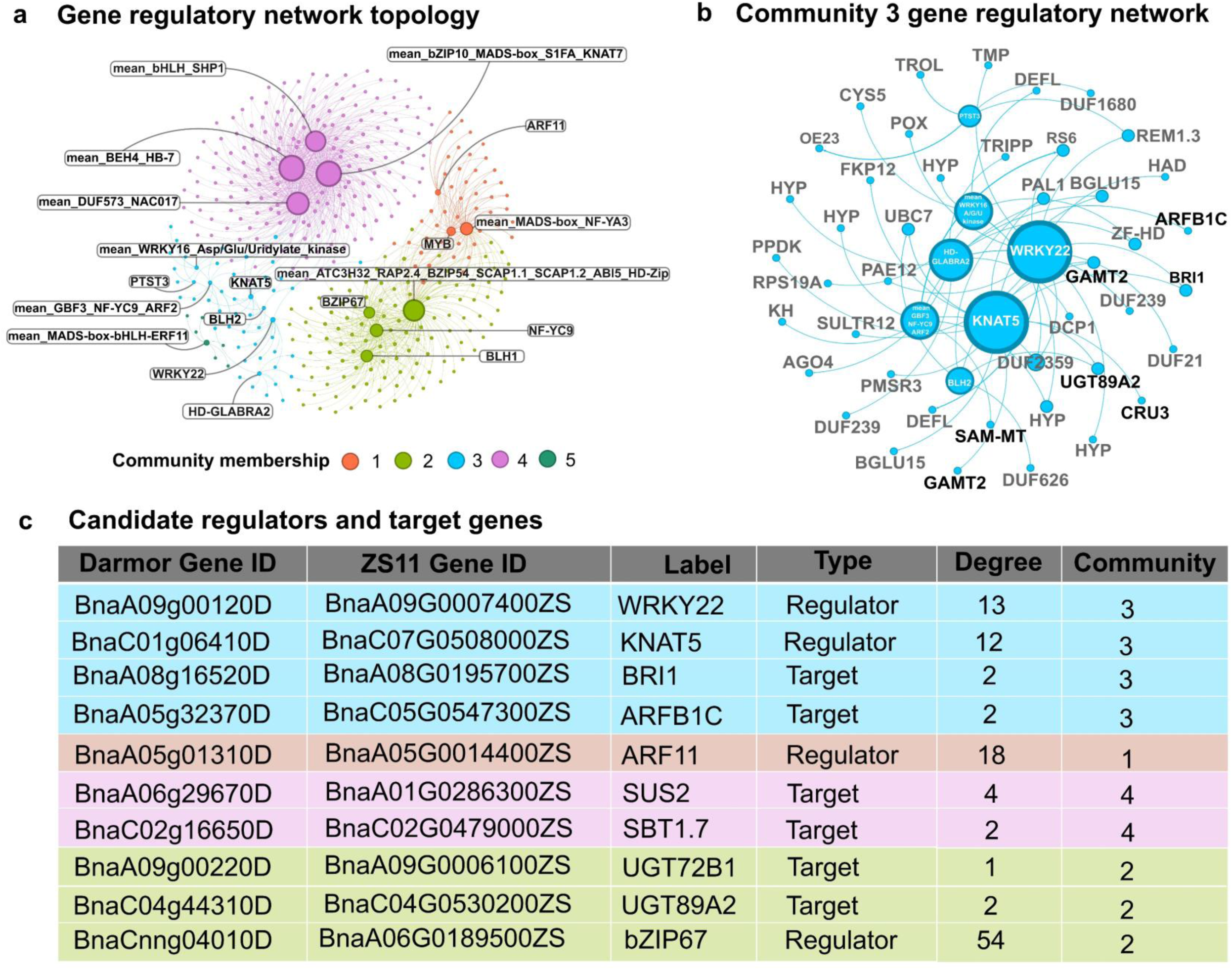
Gene regulatory network inference depicting key features and regulators involved in seed size development. (a) Network view with highlighted transcription factors in text boxes. The inferred community from the network analysis is color-coded as follows: orange (Community 1); light green (Community 2); light blue (Community 3); purple (Community 4), and dark green (Community 5). The node size is proportional to the network degree. (b) Overview of network topology in Community 3, encompassing key regulators involved in seed development. Regulators are highlighted in white, key target genes in black, and other genes belonging to the same community are in grey. (c) List of 10 influential key target genes and regulators selected based on their functional attributes and overall degree. The shading colour behind the text corresponds to the community to which the gene belongs.

To identify potential regulators, we relied on the highest degree of connectivity with the target gene. Additionally, we manually cross-checked the functional attributes of each target-gene links in each community to ensure their biological relevance. We complemented the verification step with a blast search of the *B. napus* var. Darmor gene against *B. napus* cv. ZS11 (Chen et al. 2021) and *A. thaliana* TAIR10 (Lamesch et al. 2012) genomes. This identified community 3 as the most biologically relevant, as it harbours a majority of genes involved in seed development (**Figure 6b**, **Table S6, Table S7**). The ranking of regulators in this community, based on their degree, yielded seven candidate regulators. In descending order of importance, we identified *WRKY22*, *KNAT5*, *HD-GLABRA2*, mean_*GBF3*_*NF-YC9*_*ARF2*, and mean_*WRKY16*_*A/G/U* Kinase as the regulators of this gene community. The most influential regulator, *WRKY22*, was predicted to be linked to the brassinosteroid (BR) receptor *BRASSINOSTEROID INSENSITIVE 1* (*BRI1*), a key gene involved in the maternal control of seed size in rice (Morinaka et al. 2006) and Arabidopsis (Jiang et al. 2013).

The GRNs were enriched in phytohormone pathways, including gibberellins (GAs), with *Gibberellin Methyltransferase 2* (*GAMT2*), which encodes an enzyme that catalyzes the methylation (and subsequent deactivation) of gibberellins. These plant hormones are known to regulate various developmental processes, including seed development (Varbanova et al. 2007). The second main regulator was a *KNOTTED-LIKE HOMEOBOX* gene (*KNAT5*), another Gibberellin pathway associated gene, which is assumed to regulate meristematic cells, and secondary cell wall formation (Hay et al. 2002; Rosin et al. 2003).

This analysis also identified the ARF transcription factor *ARF11*, a gene related to auxin pathway, a phytohormone that is involved in seed development.

In Community 3 we found the UDP-glycosyltransferase gene, *UGT89B1*, which catalyses the glucose conjugation of monolignols and is essential for normal cell wall development and lignification (Lin et al. 2016), likely affecting seed size (Cao et al. 2024). Moreover, two UDP-glycosyltransferase genes (*UGT72B1* and *UGT89A2*) were found in Community 2 (**Figure 6c**).

We also identified the ADP-ribosylation factor (*ARFB1C*), which has been shown to increase organ and seed size by promoting cell expansion in maize (Wang et al. 2016), and Arabidopsis (Jincheng-Yuan et al. 2015). Community 2 contains the basic *LEUCINE ZIPPER TRANSCRIPTION FACTOR* (*bZIP67*), known to regulate various seed traits in Arabidopsis, including seed size, dormancy (Bryant et al. 2019), lipid composition, and omega-3 fatty acid content (Mendes et al. 2013).

Lastly, Community 4 contains two genes of interest, *SUCROSE SYNTHASE 2* (*SUS2*) and *SUBTILASE 1.7* (*SBT1.7*) (**Figure 6c**), which are involved in cell division, starch accumulation and the control of cotyledon cell number, in cereals (Ruan et al. 2003; Li et al. 2013; Fan et al. 2019), and legumes (D’Erfurth et al. 2012).

Overall, the predicted network topology along with the biological attributes of both regulator and target genes indicated diverse pathways potentially involved in the orchestration of genes involved in seed development.

## Discussion

The agricultural relevance of seed size has driven the identification of multiple regulatory pathways, including the HAIKU pathway, the ubiquitin–proteasome system, GTPase signalling, mitogen-activated protein kinase (MAPK) cascades, and phytohormone-mediated signalling pathways, which are implicated in the control of seed size (Li and Li, 2016; Zhang et al. 2023). Endosperm growth dynamics are important determinants of seed size. The rapid growth of the endosperm by nuclear proliferation and cellular division creates the internal volume for the seed and limits embryo growth, impacting final seed size (Sundaresan, 2005; Xu and Zhang, 2023).

The polyploid crop *Brassica napus* exhibits phenotypic variation in seed size (Geng et al. 2025), yet the molecular mechanisms regulating this trait are limited. The complete transcriptomics analysis performed across four developmental embryo stages for three distinct seed tissue types enable us to confidently detect differentially expressed genes, with different gene expression between ZS11 and RJ that could play a role in the variation of seed size observed among these two genotypes, especially in the endosperm. Known genes involved in the control of seed size were detected, as the brassinosteroid (BR) receptor *BRI1* and ABI5 in the endosperm. In *A. thaliana*, *BRI1* activates the transcription factors *BRASSINAZOLE RESISTANT1* (*BZR1*) and *BRASSINAZOLE RESISTANT2* (*BZR2*) to regulate BR-responsive genes (Jiang et al. 2013). BR-deficient (*det2*) and BR-insensitive (*bri1-5*) mutants produce smaller seeds than wild-type plants, whereas exogenous BR increases seed weight (Jiang et al. 2013). *BRI1* promotes seed growth by regulating genes involved in seed size control, such as *SHORT HYPOCOTYL UNDER BLUE1*(*SHB1*), *HAIKU1* (*IKU1*) and *HAIKU2* (*IKU2*) (Jiang et al. 2013) while *ABI5* negatively regulates seed size by promoting the endosperm cellularization (Cheng et al. 2014). Co-expression analysis followed by GRN inference identified candidate genes (*WRKY22, TUBA4, TUBA5, ATARFB1C,* and *ARF17*) that were more highly expressed in ZS11 than in RJ. Notably, this analysis revealed groups of genes with similar expression patterns, suggesting a potential role in the endosperm.

The gene regulatory network enabled us to detect key regulators involved in the control of seed size. The brassinosteroid (BR) receptor *BRI1* emerged in network community 3, which activates *BZR1* that targets maternal genes such as *APETALA2*(*AP2*), *AUXIN RESPONSE FACTOR 2* (*ARF2*) and *KLUH* (*KLU*), coordinating embryo, endosperm and integument growth to positively influence seed size (Jiang et al. 2013). In community 2, we have identified a member of this gene family, *BnaA05G0014400ZS*, which encodes *ARF11,* as a potential regulator. AUXIN RESPONSE FACTORS (ARFs) are transcription factors that serve as crucial regulators of auxin-mediated gene expression (Li et al. 2023). ARFs bind to auxin responsive elements (AuxREs) in the promoters of auxin responsive genes and regulate their expression. This gene family has diverse functions, and loss-of-function mutants demonstrate that ARF transcription factors play key roles in plant development (Li et al. 2023). Although *ARF2* is the most known gene of this family for its role of controlling seed size growth in Arabidopsis (Schruff et al. 2006), it has been demonstrated that *ARF11* is positively involved in seed size in rice (Sims et al. 2021).

In addition to *BRI1*, we identified the transcription factor *WRKY22* within community 3, suggesting its potential involvement in BR signalling. To our knowledge, *WRKY22* has not yet been functionally demonstrated to regulate seed size, although other *WRKY* members have been extensively validated. In fact, the transcription factor *OsWRKY53*, has been identified in rice, as a key regulator of BR signalling, influencing leaf angle, and grain size through its interaction and phosphorylation by the *OsMAPKK4*-*OsMAPK6* cascade (Tian et al. 2017). *WRKY* genes have been found to play an important role in the regulation of seed size in other plant species, including Arabidopsis (Johnson et al. 2002; Garcia et al. 2005), soybean (Gu et al. 2017) and foxtail millet (Xiang et al. 2017). In Arabidopsis, *WRKY10*, also known as *MINISEED3*, is expressed in the developing endosperm from the 2-nuclei stage through cellularisation (Luo et al. 2005), indicating its role in endosperm development, a critical factor in determining final seed size. In addition, the expression level of the *WRKY15* is predicted to be correlated with seed size variation in both wild and cultivated soybean (Gu et al. 2017), suggesting its involvement in seed size regulation.

In foxtail millet, *LOOSE PANICLE1* (*LP1*) encodes a *WRKY* transcription factor, and loss of *LP1* function results in loose panicles and enlarged seeds (Xiang et al. 2017). This suggests that *LP1* negatively regulates grain size in foxtail millet, further highlighting the role of *WRKY* transcription factors in seed size control.

Furthermore, we also identified several candidate genes, including *KNAT5*, *ARFB1C*, *SUS2*, *SBT1.7*, *UGT72B1*, and *UGT89A2*. *KNAT5*, a KNOTTED-like homeobox transcription factor, is known to mediate meristematic development, which are regions of active cell division in both monocot and dicot (Kuijt et al. 2014). Besides, it has been observed to influence cell elongation processes in Arabidopsis (Lincoln et al. 1994; Chuck et al. 1996) and potato (Rosin et al. 2003). Further, KNOTTED1-Type Homeobox interacts with the gibberellin (GA) signalling pathway, which is known to regulate stem and leaf plant growth and development (Hay et al. 2002; Rosin et al. 2003). Meanwhile, Jia et al. (2021) concluded that *KNOX19*, a class II KNOTTED-like homeobox transcription factor from apple, is a positive regulator of the maternal seed development gene, *ABI5,* thereby, promoting organ development via ABA signalling route. Therefore, we hypothesized that *KNAT5* could interact with GA pathways that regulate plant growth as well as seed size through cell division.

Meanwhile, *BEL1* is another homeodomain transcription factor that interacts specifically with a subset of *KNOX* (*KNAT*) proteins in *Arabidopsis*, including *STM*, *KNAT1*, *KNAT2*, and *KNAT5* (Bellaoui et al. 2001). These interactions form heterodimeric complexes that regulate key developmental processes (Bellaoui et al. 2001). In the context of seed size, we hypothesize that the *BEL1*–*KNAT5* interaction influences seed size by modulating integument growth and differentiation. Since *BEL1* regulates integument initiation and development (Bencivenga et al. 2012), *KNAT5*’s specific association with *BEL1* may alter the timing and extent of integument cell proliferation and expansion, thereby affecting seed coat formation and ultimately determining final seed size.

Another candidate gene, *ARFB1C*, belonging to ADP-ribosylation factor (ARF) GTPase, is known class of genes known to be involved in regulating vesicle trafficking, thereby influencing endosperm and embryo development via cell expansion and division. (McElver et al. 2000; Gebbie et al. 2005). For instance, *ZmARF2*, an ADP-ribosylation factor expressed during early endosperm development, promotes growth in leaves, seeds, and stems by enhancing cell expansion when overexpressed in Arabidopsis (Jincheng-Yuan et al. 2015; Wang et al. 2016).

Sucrose synthase genes (SUS) have been shown to be critical for cell initiation, cell elongation, and seed development (Ruan et al. 2003; Fan et al. 2019), and its suppression in maternal seed tissues impairs fiber growth, while further suppression in the endosperm and embryo halts overall seed development (Ruan et al. 2003). Overexpression of *SUS2* in rice enhances hull size and grain weight (Fan et al. 2019) indicating SUS role in regulating cell division and starch accumulation for improved grain yield.

It is worth noting that the expression of subtilase gene *SBT1.1*, in the endosperm of *Medicago truncatula* and *Pisum sativum*, was found to be correlated with seed weight QTLs (D’Erfurth et al. 2012). Notably, mutants of *SBT1.1* show reduced seed weight and area due to decreased cotyledon cell numbers (D’Erfurth et al. 2012), implying that *SBT1.1* regulates seed size by controlling cell division.

As for UDP-glycosyltransferase, Lin et al. (2016) study revealed that UDP-glycosyltransferase 72B1 (*UGT72B1*) is crucial for normal cell wall lignification in *A. thaliana*. The *UGT72B1* mutant shows increased and ectopic lignification, arrested growth, and anthocyanin accumulation due to impaired glucose conjugation of monolignols. Therefore, *UGT72B1* plays an important role in monolignol glucosylation, which is essential for regulating lignin biosynthesis and cell wall development (Lin et al. 2016).

These candidate genes, along with the maternal seed size regulators, may collectively contribute to the complex genetic architecture underlying natural variation in *B. napus* seed size. Further functional characterization of these genes will provide valuable insights into the molecular basis of seed development.

## Conclusion

The present study used multiple approaches incorporating differential gene expression analyses, co-expression network clustering, and gene regulatory network inference to pinpoint key candidate genes, transcription factors and regulatory modules that are potential master switches for seed size development. Apart from confirming previously reported genes such as *ABI5*, and *BRI1*, new candidate genes including *KNAT5*, *ARF11*, *SUS2*, *UGT72B1*, *UGT89A2* and *SBT1.7* were also identified. Collectively, our work has provided a comprehensive insight into regulatory network involved in seed size development and advanced our understanding of genetic control of seed size in this important oilseed crop.

## Data availability

The raw RNA-seq datasets generated in this study are listed in Table S1 and available in the European Nucleotide Archive (ENA) under accession number PRJEB86494. The expression dataset is described in Woolfenden et al. (2025) and available online in the SeedORDER browser at https://seed-order-brassica.jic.ac.uk/

## Code availability

The code used in this study has been archived on GitHub (https://github.com/Yedomon/bravo_project/) and a freely accessible copy is also available on Zenodo (https://zenodo.org/records/17965416).

